# Short gene overlaps in bacterial genomes: evolutionary dynamics and functional associations

**DOI:** 10.64898/2026.09.10.750558

**Authors:** Arseniy Yu. Sukhodolsky, Anna D. Kaznadzey, Mikhail S. Gelfand

## Abstract

**Background:** Short overlaps of adjacent genes are widespread in bacterial genomes and have been proposed to contribute to coordinated gene expression through mechanisms such as translational coupling and ribosome re-initiation. However, their evolutionary dynamics and functional associations have been studied either in relatively small datasets or for individual taxonomic groups. Here, we performed a large-scale comparative analysis of bacterial gene overlaps to characterize their prevalence, structural diversity, evolutionary dynamics, and functional associations. We surveyed 3,998 representative bacterial genomes for short overlaps and analyzed a curated set of orthologous gene-pair clusters to reconstruct the evolutionary history of overlap gain and loss using ancestral-state inference.

**Results:** We confirmed that short overlaps are dominated by a small number of canonical configurations, while a substantial subset of orthologous clusters contained multiple overlap lengths or sequence motifs, indicating evolutionary flexibility. Reconstruction of ancestral states yielded repeated gains and losses of overlap within the same orthologous gene-pair lineages. Among clusters in which both states were sufficiently represented, we found no overall bias toward either overlap gain or loss. Finally, analyses of functional associations found no consistent enrichment of overlapping gene pairs among genes encoding interacting protein subunits, compared to closely spaced non-overlapping gene pairs. In contrast, metabolic linkage showed a modest positive association with overlap, although the magnitude of this association depended on the distance threshold used to define non-overlapping neighbors.

**Conclusions:** These results identify short bacterial gene overlaps as evolutionarily labile features of gene organization rather than strongly conserved genomic states and emphasize the importance of distinguishing overlap from close genomic proximity when considering their functional significance.

## Background

Gene overlap refers to sharing of one or more nucleotides between coding sequences. This phenomenon was first discovered in the genome of bacteriophage φX174, a small single-stranded DNA virus (5386 bp) that infects *Escherichia coli* [1, 2]. It was previously thought to occur mainly in viral genomes, but it was later found that gene overlaps occur in genomes of all domains, from prokaryotes to eukaryotes [3].

Gene overlap in prokaryotes refers to two coding sequences (CDSs) that share one or more nucleotides [3]. Three different overlap types are fundamentally possible. Unidirectional overlap is observed between genes encoded within a single strand. For genes located on opposite strands, two alternative configurations exist. Divergent configuration refers to a case when start codons of the genes overlap and convergent refers to a case when stop codons overlap [3]. Prokaryotic and viral genomes exhibit a significant prevalence of unidirectional topological arrangements with the prevalence of short overlaps, usually 1 or 4 bp (base pairs) [4].

Although gene overlaps have been proposed to have diverse biological functions, their precise role and evolution remain incompletely understood. Overlapping gene organization may facilitate fusion of functionally related genes [5] and contribute to regulation of translation [6]. Functionally, gene overlap may be an adaptive strategy, as it fine-tunes regulation via translational coupling and ribosome re-initiation [3, 6, 7]. It was shown for *H. volcanii* and *E. coli*, that overlapping and closely spaced unidirectional gene pairs are enriched in genes encoding subunits of heteromeric complexes [8].

It is believed that overlapping genes in prokaryotes on average evolve slower than non-overlapping genes. At the same time, overlaps themselves are much less conserved [3]. Both the intergenic distance for neighboring genes and the length of the overlap can vary even between evolutionary close species [9].

Although several mechanisms of overlap formation have been proposed, the evolutionary dynamics of short overlaps remain poorly characterized across bacteria. In particular, it remains unclear how frequently overlaps are gained and lost and whether these transitions show a general directional bias. Understanding these dynamics is also relevant for the rational design of engineered overlapping genes used in synthetic biology [10].

Our goal was to comprehensively characterize the prevalence, diversity, and evolutionary dynamics of short bacterial gene overlaps (1–10 bp). Specifically, we aimed to (i) quantify the prevalence and structural characteristics of short overlaps across bacterial genomes, (ii) estimate the rates of overlap gain and loss during evolution, and (iii) investigate whether overlap occurrence is associated with functional relationships between neighboring genes. We found that although short overlaps are concentrated in a limited number of canonical configurations, overlap itself represents an evolutionarily dynamic state that can repeatedly arise and disappear in orthologous groups of gene pairs. Analyses of functional associations revealed no consistent enrichment of overlapping gene pairs among genes encoding interacting protein subunits, suggesting that protein complex membership alone does not explain the evolutionary distribution of short overlaps.

## Methods

### Data selection

Bacterial genomes were obtained from the Refseq database [11]. We selected genomes which were marked as reference or representative, one for each species. Short gene overlaps, from 1 to 10 base pairs, were selected, resulting in 15066614 genes with 1783435 detected short overlaps from 3998 genomes.

### Initial grouping

Firstly, we identified pairs of overlapping genes. The eggNOG-mapper 5.0 [12] was used to assign each gene to a certain family. Each pair of families formed a group of respective gene pairs. To analyze cases where genes are non-overlapping neighbors, we also added to our groups pairs of genes which were located within a short distance (<100 bp) from each other and belonged to respective EggNOG families. Due to limitations of the EggNOG database, the dataset was reduced to 1350 organisms for this part of the study.

### Clustering

In order to narrow down the differences between genes within our groups and also to exclude misannotations, we further categorized the groups into clusters using MMseqs2 [13]. We used the greedy set cover algorithm, with minimum sequence identity = 0.3 and –alignment-mode 3. The resulting number of clusters was 3554.

### Data purification and misannotation management

To align overlapping regions within clusters, we used the Clustal Omega tool [14], selecting clusters of over 20 pairs in size (3139 clusters containing 112450 gene pairs in total). We consider a start as misannotated, if the start codon is not in the same location as most start codons of other genes of the alignment, but at the ‘correct’ location (which is no more than 20 bp away), there is a possible start codon.

**Figure 1.**
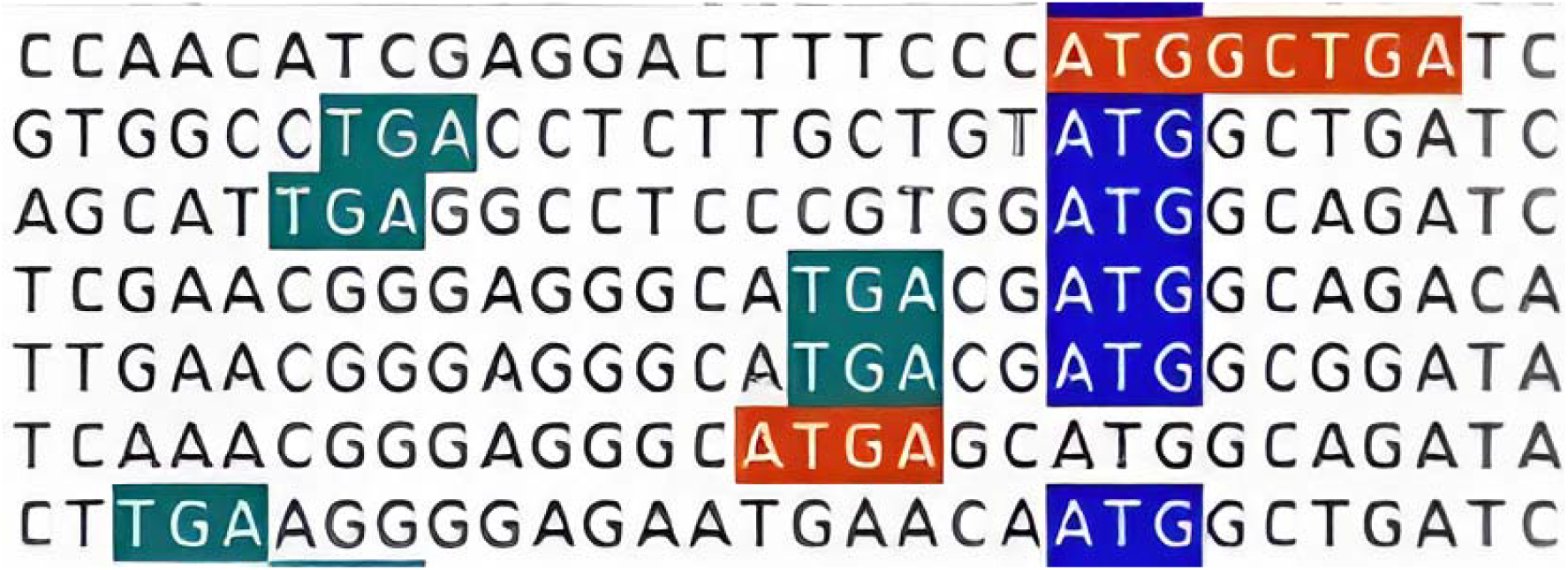
An example of overlap misannotation. Red refers to annotated overlap, green refers to the annotated stop codon of the first gene, blue refers to the annotated start codon of the second gene. While the top overlap, ATGGCTGA, is real, the bottom one, ATGA, results from a likely misannotated, conserved start codon ATG.

Clusters in which start codons were overall poorly aligned (i.e. less than 90% of aligned genes have start codons at the same location) were excluded from further investigation. We identified 199 misannotations based on this rule. The resulting dataset contained 484 clusters with 20 or more gene pairs each.

### Overlap variability

To further characterize the variability of overlap within orthologous clusters, we analyzed the distribution of overlap lengths and sequence motifs across clusters. For each cluster, we identified all overlap lengths and motifs and asked how often clusters are fixed for a single type versus harbouring multiple co-occurring variants.

### Reconstruction of ancestral states

In order to estimate the evolution tendencies and rate of the gene overlapping we used BayesTraits [15] for evolutionary reconstructions. Since our model assumes only two gene pair states (overlapping and non-overlapping), we used a two-state model for evolution. *q*_gain_ and *q*_loss_ represent the instantaneous transition rates from non-overlapping to overlapping and from overlapping to non-overlapping states, respectively [16]. AnnoTree was used as the reference bacterial phylogenetic tree [17]. To predict ancestral states we pruned that tree for each cluster using the ete3 python module [18]. For each orthologous cluster, gain and loss rates were estimated as posterior medians from a continuous-time Markov model fitted in BayesTraits. As options we used the MultiState mode and the MCMC approach. All MCMC parameters and prior distributions were left at the BayesTraits default settings.

The initial dataset included every cluster containing at least one overlapping gene pair. However, clusters with a strong imbalance between overlapping and non-overlapping states had to be excluded from the rate analysis. When one state is represented by only a few extant observations, the model cannot reliably distinguish between a recent origin of that rare state and a high transition rate away from it. Consequently, ancestral-state estimates and transition rates become poorly constrained for such clusters. Therefore, we restricted this analysis to clusters in which at least 20% of gene pairs are overlapping and at least 20% are non-overlapping, yielding 164 clusters.

### Identification of subunits

To determine whether the tendency to form overlaps is associated with the members of a gene pair encoding protein subunits, we expanded our dataset to include clusters that did not contain directly overlapping genes but included genes related to those with detected short overlaps and located at small intergenic distances. The resulting dataset comprised 257 subunit-associated and 601 non-subunit clusters.

Since our clusters were compiled based on gene pair orthology, we defined a subunit cluster as one in which at least one gene pair has a high confirmed experimental interaction level in the STRING database [19] (experimental > 700) and their annotation included one of the key words (‘subunit’, ‘component’, ‘chain’, ‘domain’, ‘fragment’, ‘complex’). We selected this cutoff to be reasonably confident in the reliability of the data regarding subunit composition. We then divided our clusters into subunit and non-subunit clusters and measured the overlap fraction for each group.

### Genome-wide functional association analysis

To assess whether gene overlap is associated with functional relationships between neighboring genes, we extended the analysis beyond the curated orthologous clusters to all neighboring gene pairs separated by no more than a specified intergenic-distance threshold (20–500 bp). For each threshold, gene pairs were classified according to the overlap status and functional relationship, and contingency tables were analyzed using Fisher’s exact test. For protein-complex association, a gene pair was considered subunit-linked if both genes encoded subunits of the same protein complex according to the STRING database [19], using only experimentally supported interactions (experimental score > 700) such that their annotation included one of the key words (‘subunit’, ‘component’, ‘chain’, ‘domain’, ‘fragment’, ‘complex’). This criterion was identical to that used for the orthologous-cluster analysis, except that it was applied directly to all neighboring gene pairs rather than to orthologous clusters.

To identify metabolically related gene pairs, all proteins were first annotated using KofamScan [20]. Each gene was assigned to KEGG Orthology (KO) groups, which were then mapped to KEGG metabolic reactions. Two neighboring genes were considered metabolically linked if at least one reaction associated with the first gene and one reaction associated with the second gene shared a common compound that was not part of a predefined set of ubiquitous (“currency”) metabolites. The following ubiquitous (“currency”) metabolites were excluded from consideration: H2O (C00001), H+ (C00080), ATP (C00002), ADP (C00008), AMP (C00020), NAD+ (C00003), NADH (C00004), NADPH (C00005), NADP+ (C00006), orthophosphate (C00009), and diphosphate (C00013). This procedure identifies neighboring genes participating in consecutive or otherwise directly connected metabolic reactions while excluding links mediated solely by common cofactors and other highly connected metabolites.

### AI Assistance

During the preparation of this manuscript, the authors used OpenAI ChatGPT (GPT-5.6 Sol) for assistance with language editing, manuscript organization, code development and debugging, statistical analysis, and figure preparation. All AI-assisted outputs were critically reviewed, verified, and, where necessary, modified by the authors. The authors take full responsibility for the content of this publication.

## Results

### Overlap statistics

Overall, 15066614 genes with 1783435 detected short overlaps from 3998 genomes were analyzed in this part of the study. We collected statistics on overlap types, identifying unidirectional and convergent overlaps.

**Figure 2.**
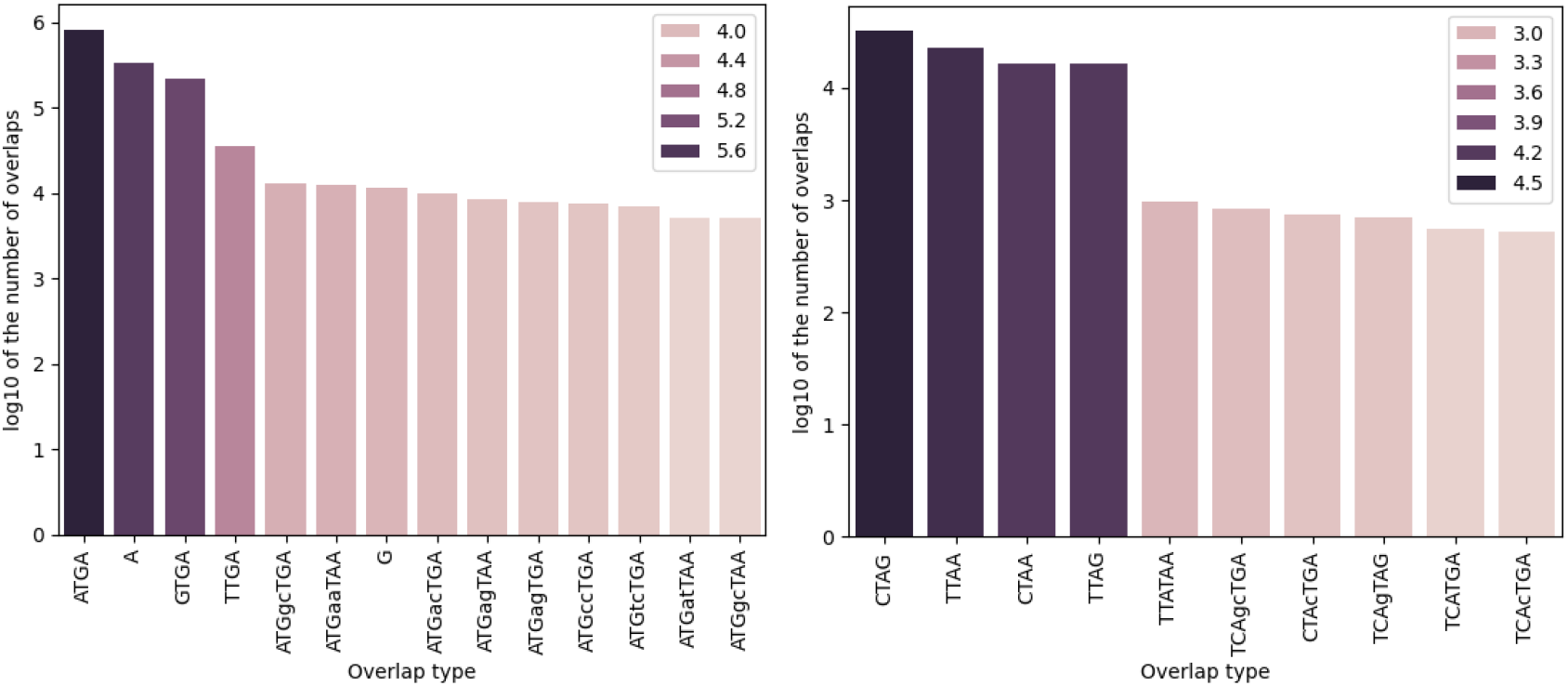
Distribution of overlap types: unidirectional and convergent cases. Nucleotides corresponding to start and/or stop codons are highlighted in capital letters.

Unidirectional overlaps are much more common than convergent and divergent ones. They constitute 92.9% of all considered overlaps, the most common length being 1 and 4 bp, although longer overlaps have been observed as well. The most frequent types were ATGA (24% of all unidirectional overlaps), A(10%) and GTGA (3.8%). In the case of divergent and convergent overlaps, we give the overlap type as it appears on a reference strand. The most common convergent types were CTAG (28% of all convergent overlaps), TTAA (14%), CTAA (7%), and TTAG (7%), the latter two representing the same type of overlap, dependent on which gene was in the reference strand. They constitute 6.2% of all considered overlaps. For example, for the overlap CTAG in the convergent case, the stop codon for the gene on the forward strand is TAG, as it is for the gene on the reverse strand. Only 0.9% of overlapping gene pairs had a divergent overlap configuration, of which 73% were of the AT type, corresponding to a start–start overlap in which the ATG start codons of both genes overlap.

### Overlap variability within orthologous clusters

The analyses presented further were performed on a more stringently filtered subset of orthologous clusters rather than on the complete collection of overlaps described above. To ensure reliable comparisons and ancestral-state reconstruction, only well-supported orthologous clusters passing all quality-control criteria were retained for downstream analyses.

**Figure 3.**
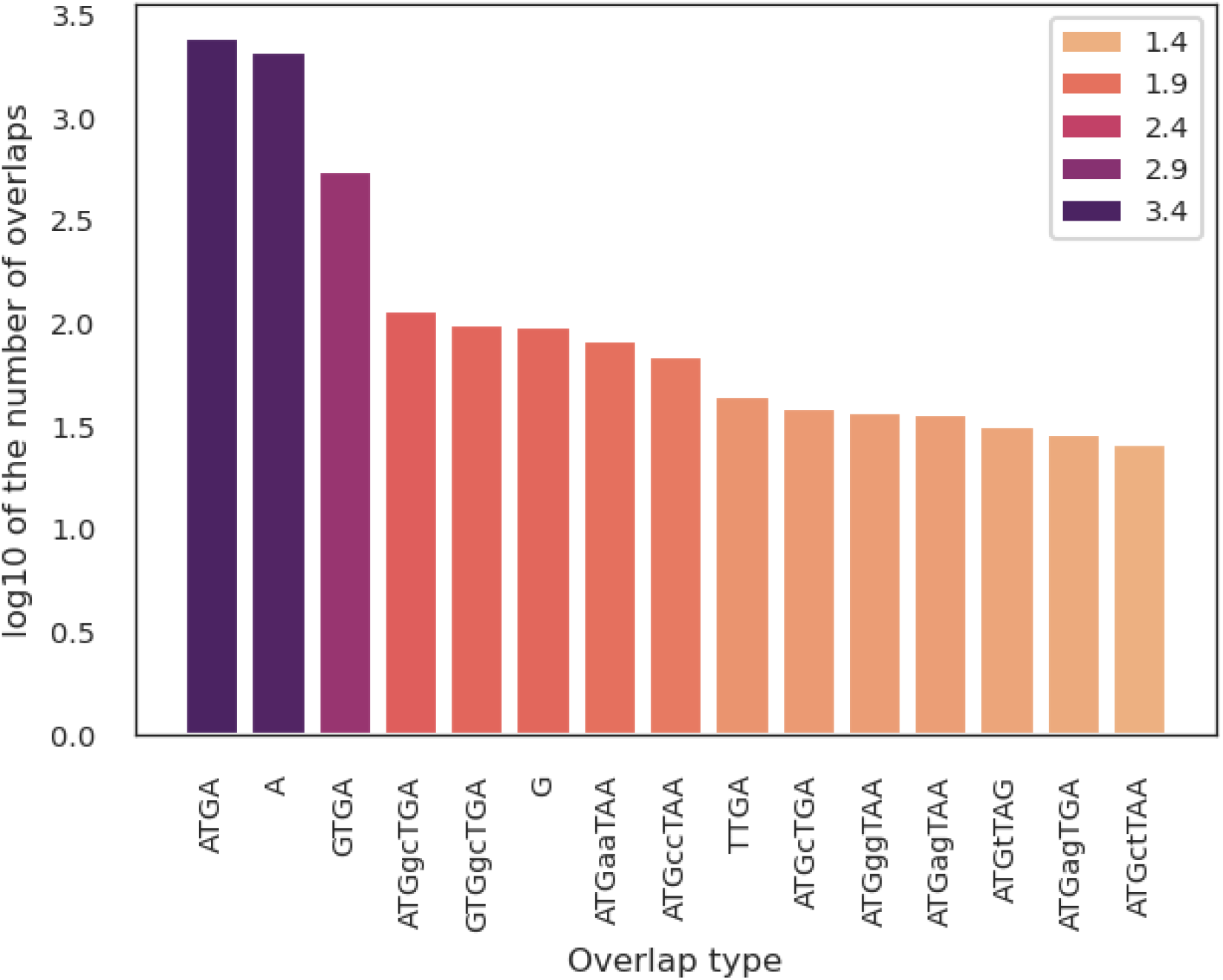
Distribution of overlap types in orthologous clusters: unidirectional case. Nucleotides corresponding to start and/or stop codons are highlighted in capital letters.

**Figure 4.**
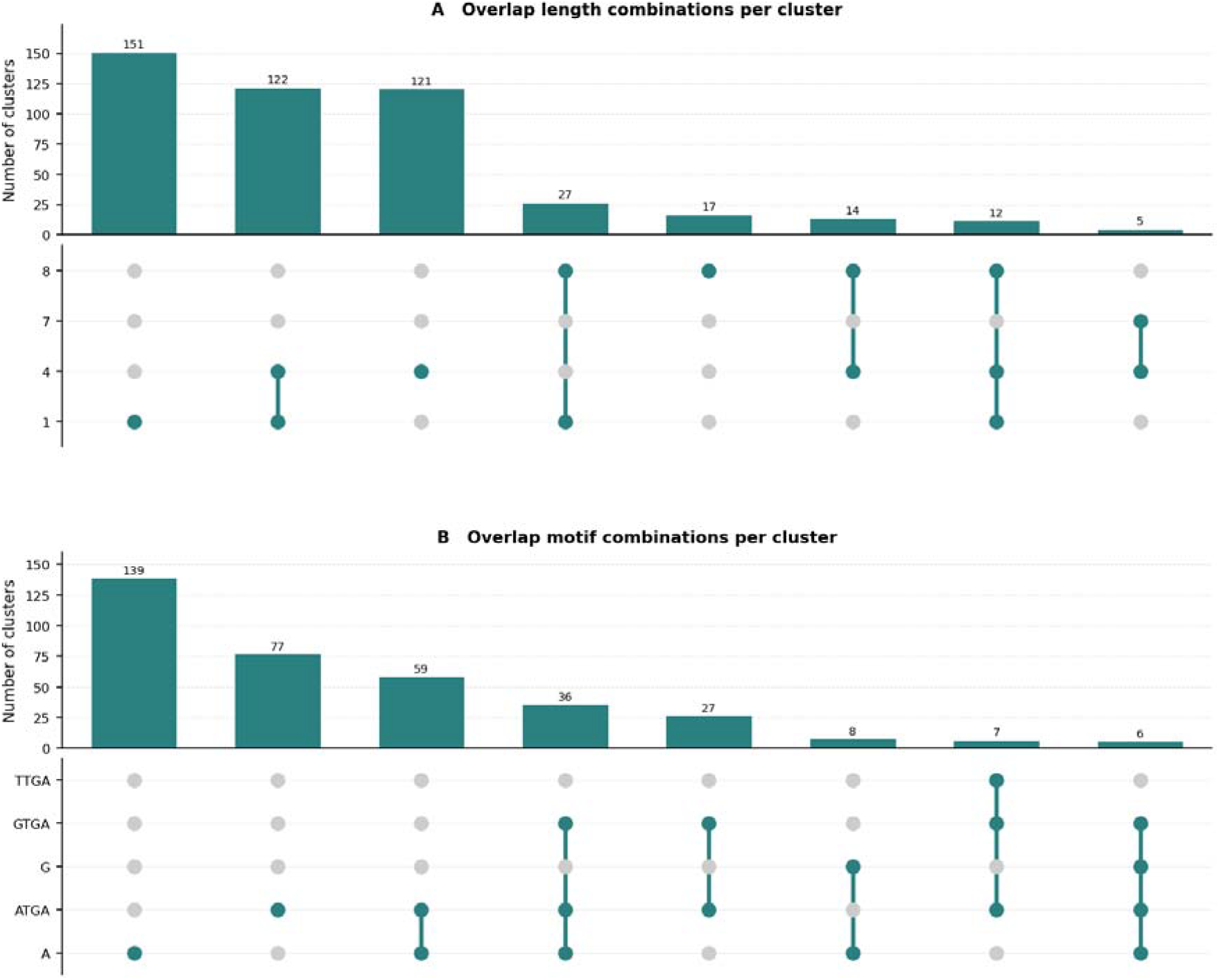
Overlap length and motif diversity in orthologous clusters. UpSet plots show the number of clusters harbouring each combination of overlap lengths (A) or motifs (B); only the top combinations by frequency are shown. Single-element bars correspond to clusters fixed for one length or motif; multi-element intersections indicate clusters where two or more variants co-occur.

The majority of clusters were characterized by a single overlap length. The most common categories were clusters containing only 1 bp overlaps (151 clusters), followed by clusters containing only 4 bp overlaps (121 clusters). Among clusters with multiple overlap lengths, the 1+4 bp combination was the most frequent (122 clusters), whereas combinations involving 1+8 bp (27 clusters), 4+8 bp (17 clusters), or three overlap lengths (14 and 12 clusters for the most common combinations) were less frequent. Thus, although many orthologous clusters retain a single characteristic overlap length, coexistence of several overlap lengths within the same cluster is also common, indicating substantial evolutionary variability of overlap architecture within orthologous gene pairs.

A similar pattern was observed at the motif level. Clusters containing only the 1 bp motif A were the most frequent (139 clusters), followed by clusters containing only ATGA (77 clusters). The most common mixed motif category was A+ATGA (59 clusters), while more complex combinations involving three or four motifs were progressively less frequent. Rare combinations involving motifs such as G, GTGA, and TTGA were also present. Overall, the motif-level distribution suggests that although a few overlap configurations predominate, multiple alternative overlap motifs can occur within the same orthologous cluster.

### Reconstitution of ancestral states

Reconstruction of ancestral states and calculation of median predicted overlap gain and loss rates (*q*_gain_, non-overlapping → overlapping, and *q*_loss_, overlapping → non-overlapping) revealed that overlap configuration is evolutionarily labile. Across many orthologous clusters, overlaps were repeatedly gained and lost, rather than being maintained as stable ancestral features. Representative examples illustrating these evolutionary patterns are shown in Figures 5-7. The reconstructed histories illustrate several characteristic evolutionary scenarios.

**Figure 5.**
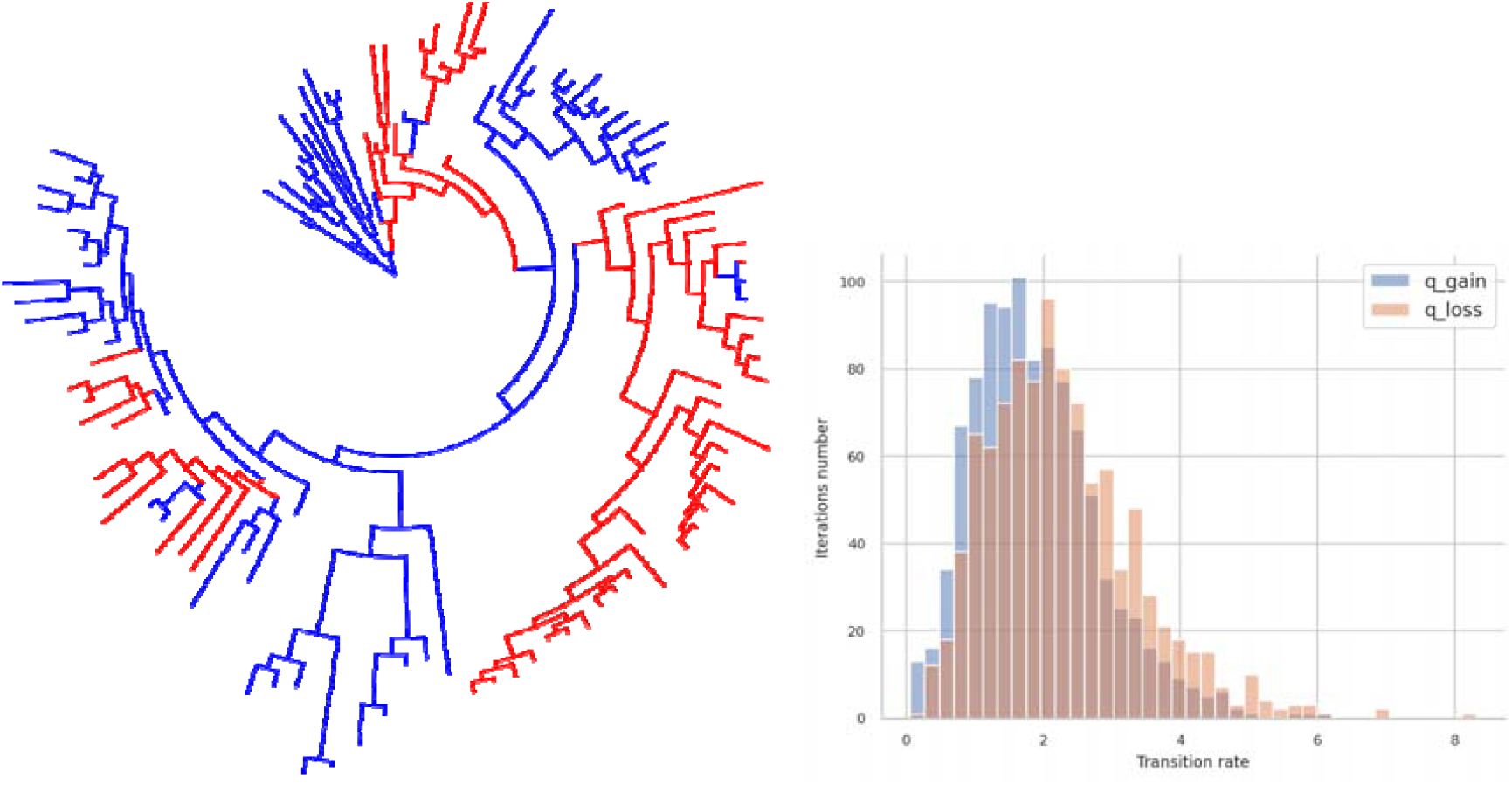
Ancestral-state reconstruction for ATP synthase subunits A and delta with balanced gain and loss rates. q_gain_ = 1.8, q_loss_ = 2.1. Red branches indicate predicted overlapping states, blue branches indicate predicted non-overlapping states, and gray branches indicate ambiguous predictions (posterior probability 45–55%). The posterior distributions of transition rates are shown at the right. Both states are well represented across the phylogeny, reflecting multiple inferred transitions between overlapping and non-overlapping states.

In the example shown in Figure 5, both overlapping and non-overlapping gene pairs are well represented across the phylogeny. For this orthologous cluster, the estimated median transition rates are similar (*q*_gain_ = 1.8, *q*_loss_ = 2.1), resulting in repeated transitions between overlapping and non-overlapping states during evolution. Accordingly, ancestral state reconstruction indicates that both states were likely present at different points in the evolutionary history of this gene family.

The second example is characterized by a substantially higher rate of overlap loss than gain (*q*_gain_ = 2.8, *q*_loss_ = 7.9). Accordingly, the reconstruction predicts that non-overlapping states predominate throughout the evolutionary history of this orthologous cluster (Figure 6). Nevertheless, the reconstructed history also includes multiple transitions in both directions, indicating that overlap can be regained after being lost.

**Figure 6.**
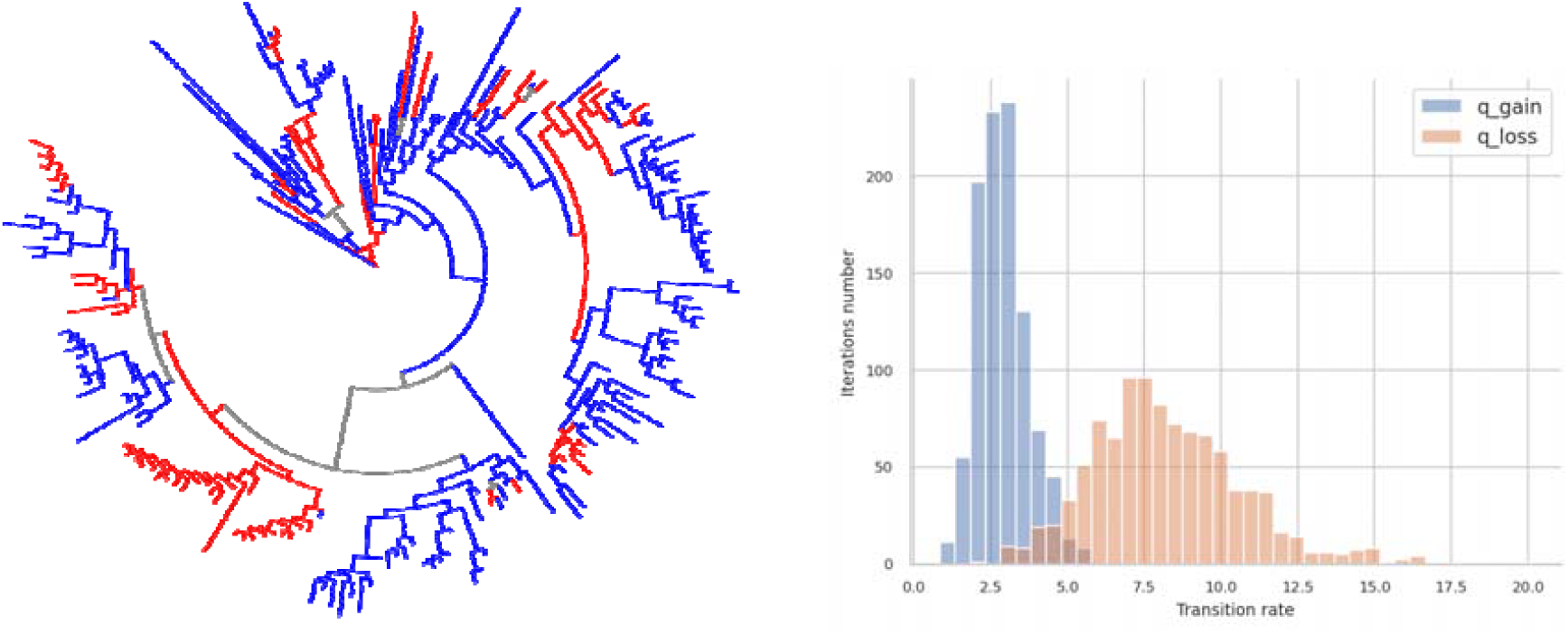
Ancestral-state reconstruction for succinyl-CoA ligase ADP-forming subunits alpha and beta with predominant overlap loss. q_gain_ = 2.8, q_loss_ = 7.9. Notation as in Figure 5. Despite the higher loss rate, overlapping states recur multiple times across the phylogeny, illustrating the dynamic nature of overlap evolution even in clusters where one state is disfavoured.

Nevertheless, some orthologous clusters are characterized by a predominance of overlapping states, an example in Figure 7. Although overlap is inferred to be the prevailing state throughout much of the phylogeny, the estimated transition rates (*q*_gain_ = 2.6, *q*_loss_ = 4.0) again indicate that transitions between overlapping and non-overlapping states occur repeatedly, even within closely related clades.

**Figure 7.**
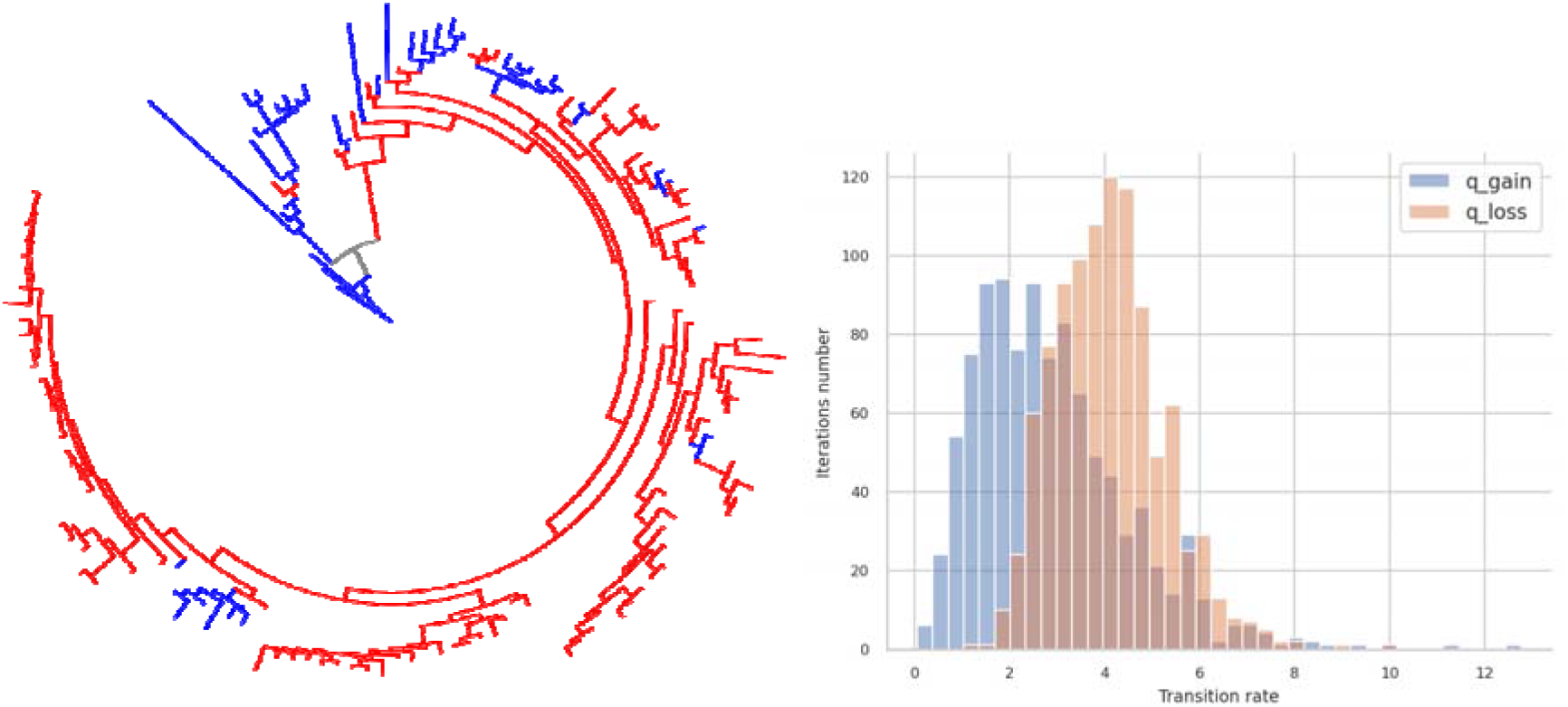
Ancestral-state reconstruction for succinyl-CoA synthetase subunits beta and alpha with high overlap prevalence. qgain = 2.6, qloss = 4.0. Notation as in Figure 5. Despite the predominance of overlapping states, transitions between states occur repeatedly even among closely related lineages, further illustrating the dynamic nature of overlap evolution.

**Figure 8.**
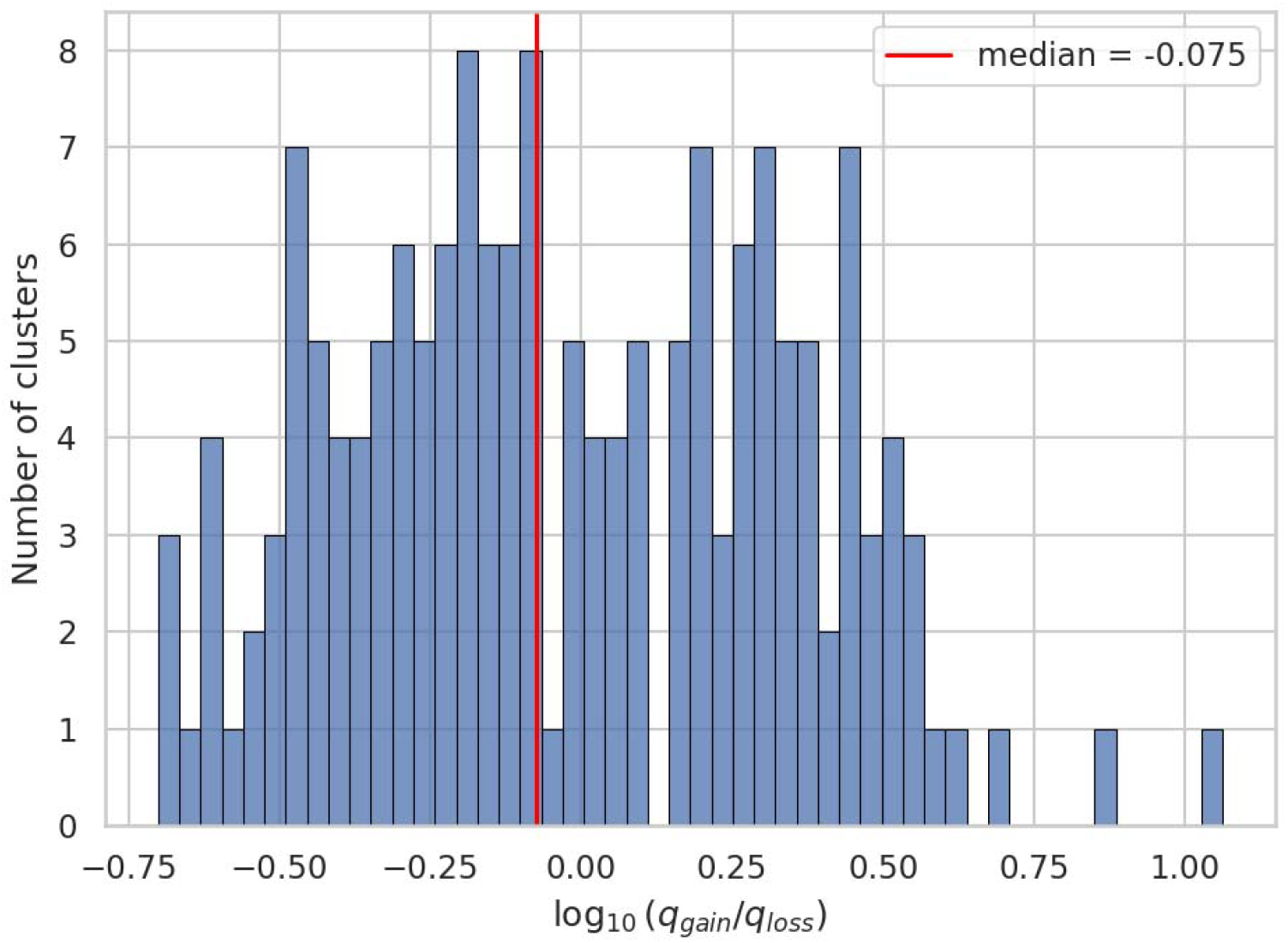
Distribution of estimated log_10_(*q*_gain_/*q*_loss_) values across orthologous clusters. Only clusters in which overlap was present in 20–80% of members were included in the analysis. Positive values indicate a higher estimated rate of overlap gain than loss, whereas negative values indicate the opposite. The vertical dashed line marks equal gain and loss rates, log_10_(*q*_gain_/*q*_loss_)=0.

Although these three clusters differ in the relative prevalence of overlapping and non-overlapping states, all show evidence of transitions in both directions. We therefore asked whether overlap gain or loss tends to predominate across the dataset. For that, we next examined the distribution of transition-rate estimates across all clusters in which both overlapping and non-overlapping states were sufficiently represented (see Methods).

The median log-ratio of posterior median gain to loss rates was −0.075 across these clusters. A two-sided Wilcoxon signed-rank test found no significant deviation of the median log_10_(*q*_gain_/*q*_loss_) from zero (*W* = 6,422, *p* = 0.49), suggesting that, among clusters with sufficiently mixed overlap states, gain and loss rates are broadly balanced. We compared transition rates between clusters whose gene pairs encode protein subunits and those that do not (Figure 9).

**Figure 9.**
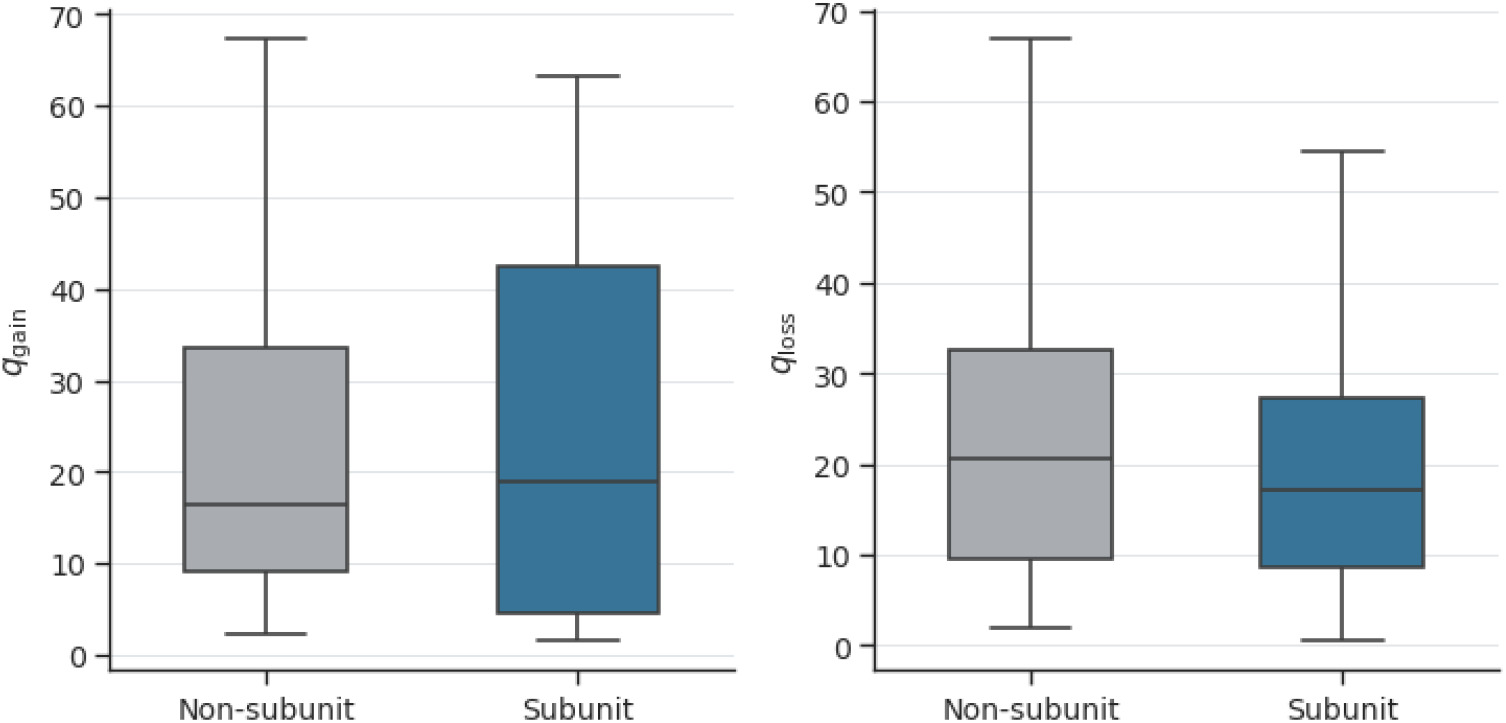
Overlap gain and loss rates in gene-pair clusters with substantial representation of both states. Distributions of cluster-level posterior median estimates of the overlap-gain rate, *q*_gain_ (left), and overlap-loss rate, *q*_loss_ (right), are shown for non-subunit (*n*=111) and experimentally supported subunit-associated (*n*=53) clusters. Only clusters in which overlapping and non-overlapping gene pairs each represented at least 20% of observations were included. Boxes indicate the median and interquartile range; whiskers extend to 1.5 times the interquartile range. Outliers are omitted for clarity.

**Figure 10.**
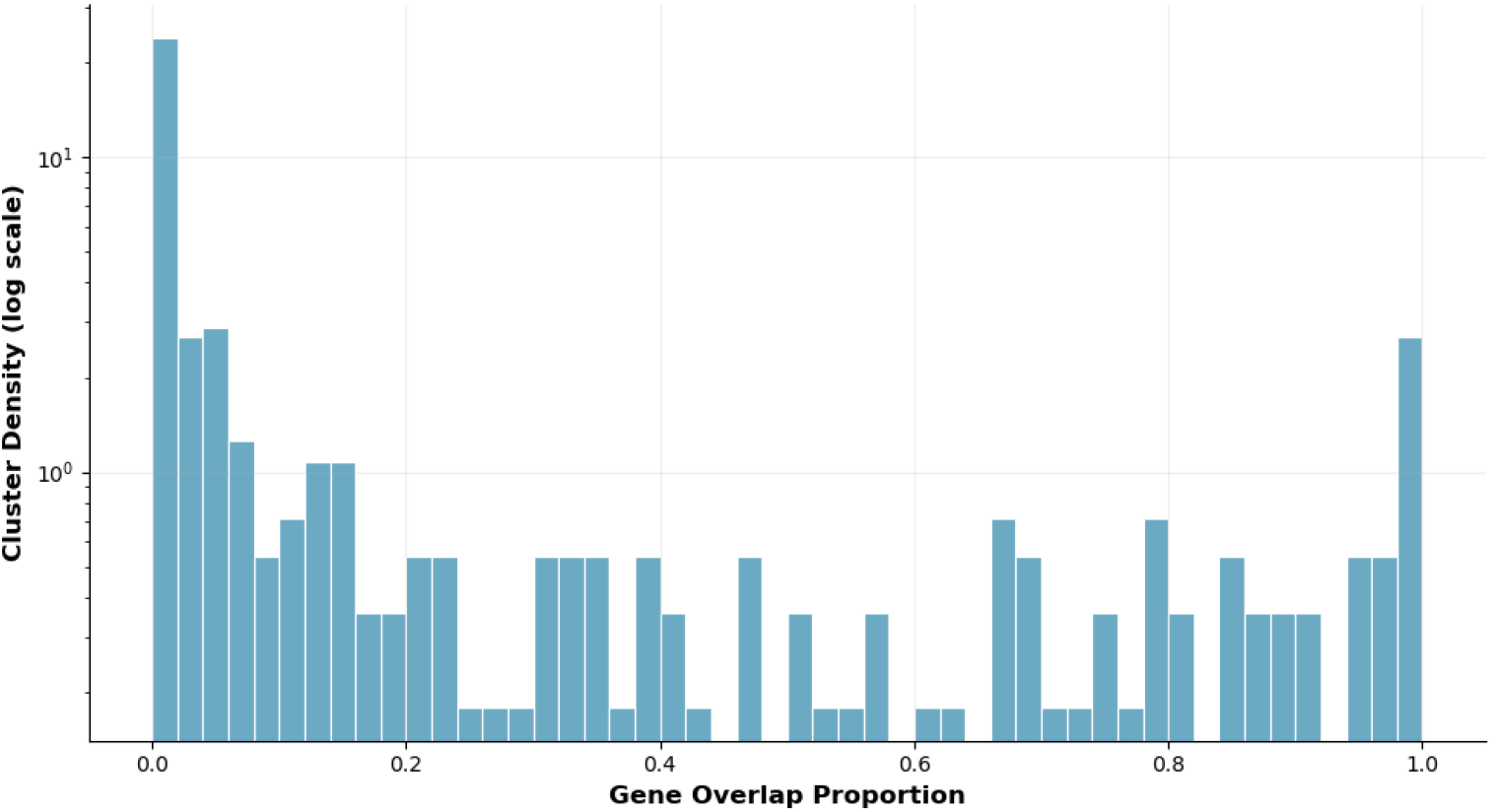
Distribution of gene-overlap proportions among orthologous clusters containing protein-complex subunits. For each cluster, the overlap proportion was calculated as the fraction of gene pairs displaying a short overlap among all gene pairs assigned to that cluster.

**Figure 11.**
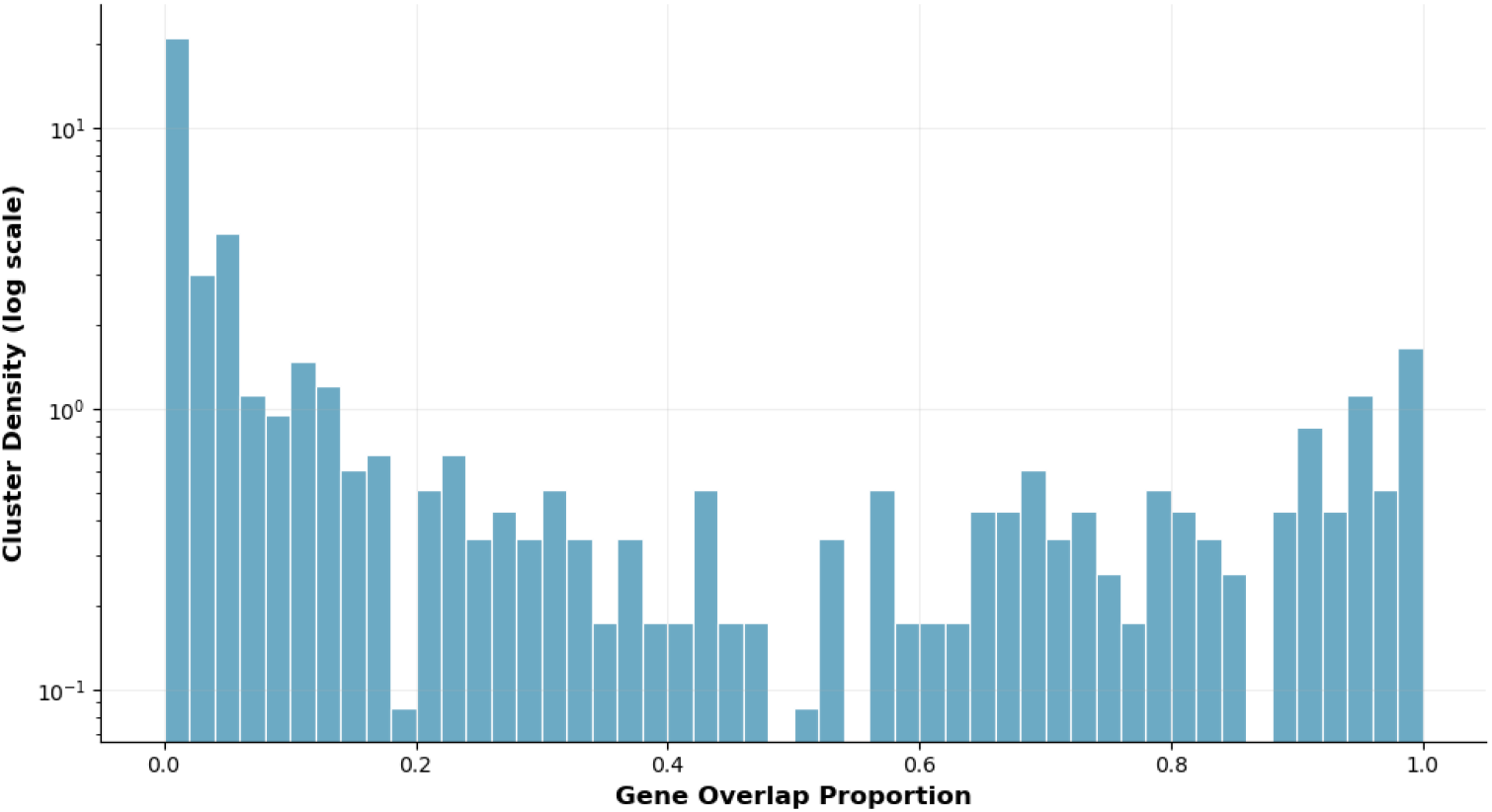
Distribution of gene overlap proportions across non-subunit clusters. For each cluster, the overlap proportion was calculated as the fraction of gene pairs displaying a short overlap among all gene pairs assigned to that cluster.

**Figure 12.**
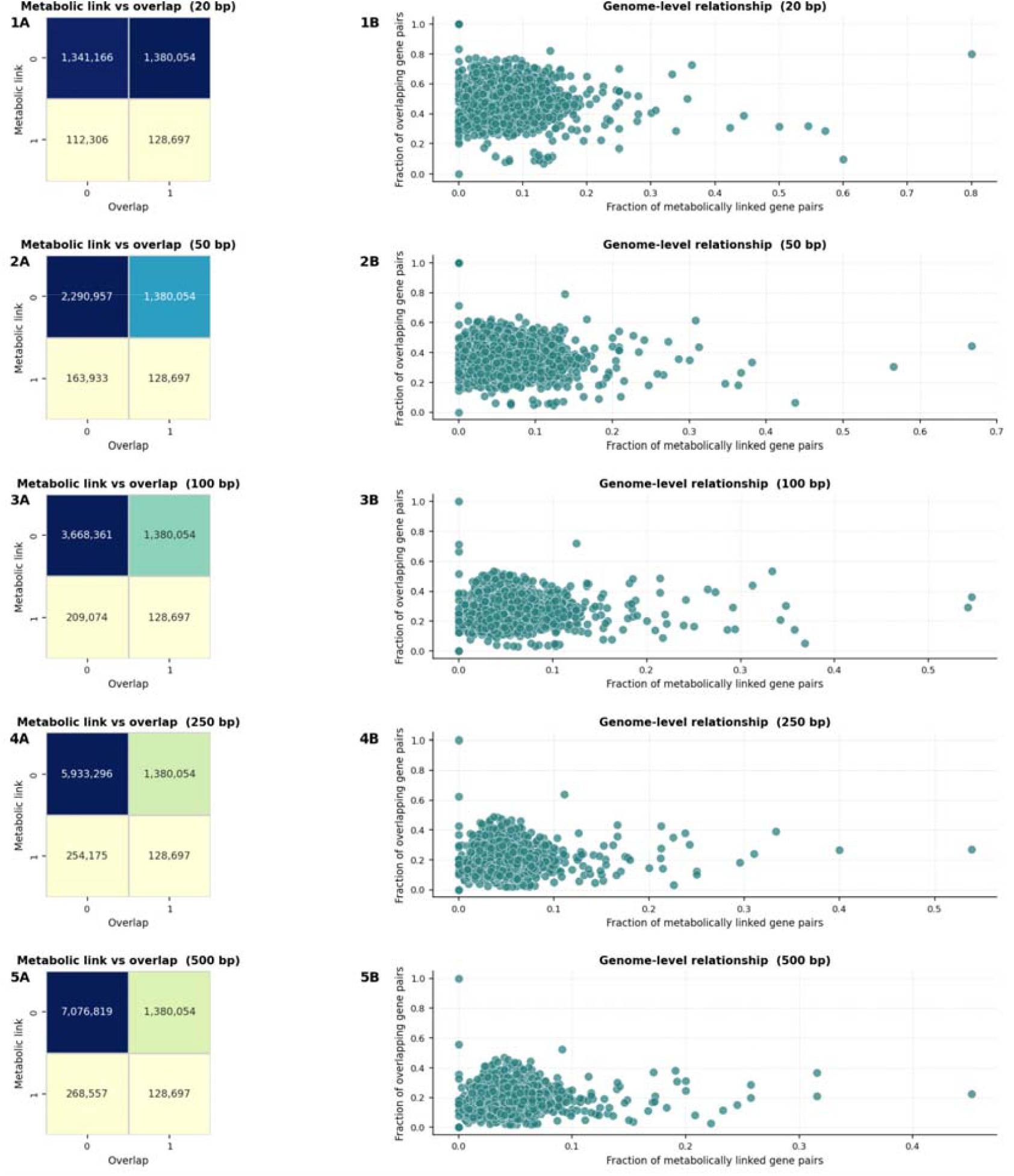
Substrate adjacency and gene-pair overlap across distance thresholds. For each threshold (20–500 bp), gene pairs were classified by whether the genes participate in metabolically linked reactions sharing at least one non-currency compound (metabolic link: no/yes) and whether they overlap (overlap: no/yes). (A) Contingency tables show the number of gene pairs in each category. (B) Each point represents one genome; axes show the fraction of gene pairs with metabolic linkage and the fraction with overlap, respectively.

**Figure 13.**
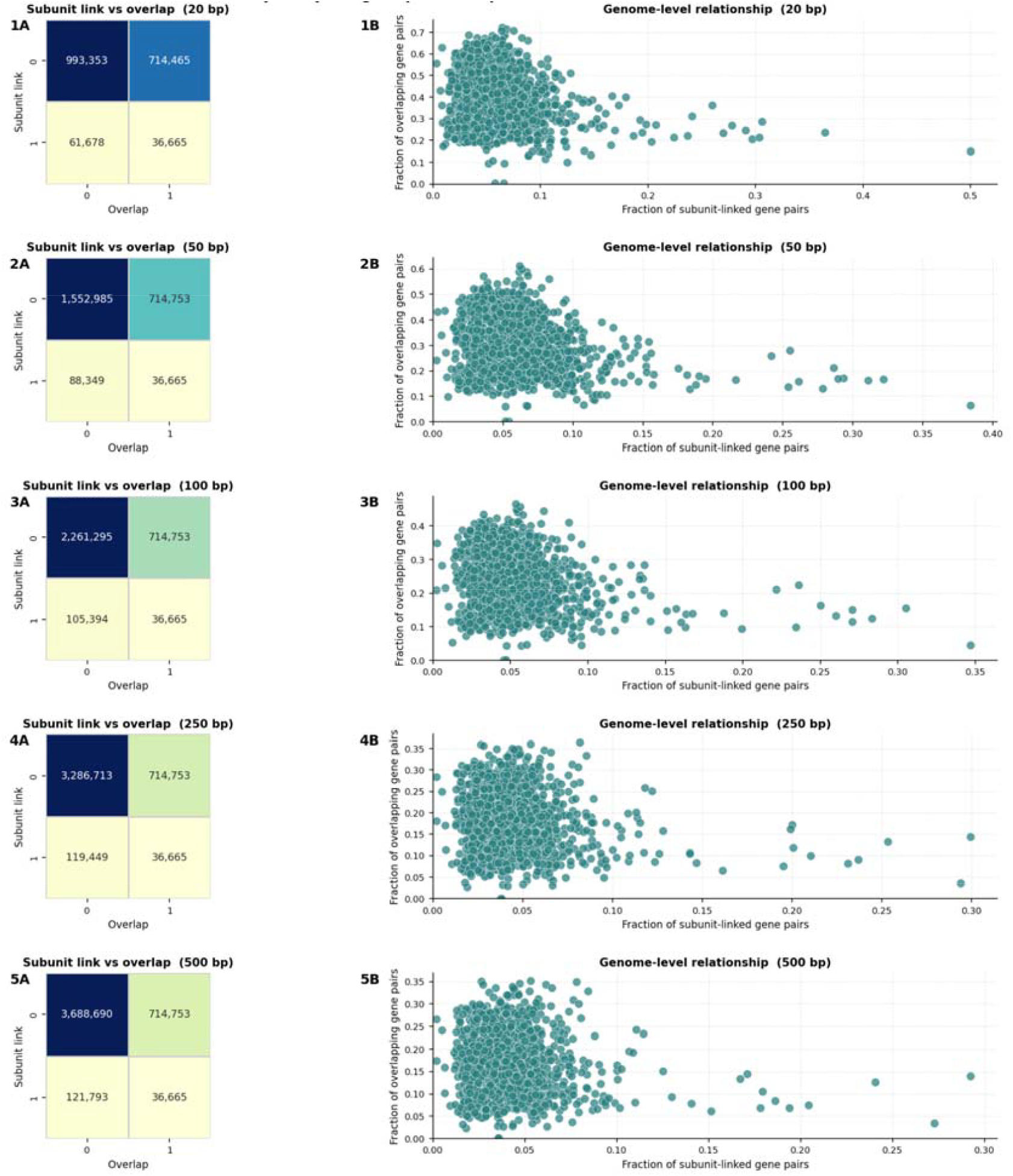
Association between subunit linkage and gene overlap across distance thresholds. Gene pairs were classified by whether both genes encode putative subunits of the same protein complex (subunit link: 0/1), as inferred from STRING experimental interaction scores (≥ 700), and whether they overlap (overlap: 0/1). For each threshold (20–500 bp), (A) contingency tables show the number of gene pairs in each category; (B) each point represents one genome, with axes showing the fraction of subunit-linked gene pairs and the fraction of overlapping gene pairs, respectively.

The median overlap fractions were nearly identical between the two groups (0.470 and 0.468, respectively). Subunit-associated clusters showed a slightly higher median overlap-gain rate than non-subunit clusters (18.95 versus 16.46) and a slightly lower median overlap-loss rate (17.14 versus 20.60). However, the within-cluster balance between gain and loss rates did not differ significantly between subunit-associated and non-subunit clusters.

### Association of gene overlap with subunits of protein complexes

We measured the proportion of overlapping pairs within orthologous clusters and compared the average overlap ratios between clusters whose gene pairs encode protein subunits and those that do not.

In clusters where gene pairs are experimentally confirmed to form subunits, the average proportion of overlapping pairs is 0.24 ± 0.34, whereas in clusters where they do not, it is 0.22 ± 0.32.

Given these results, we considered the possibility that they might be influenced by the composition of our curated orthologous-cluster dataset. Although some clusters do not contain overlapping genes, they consist of orthologs of genes that overlap in other genomes. Consequently, the dataset is inherently enriched for gene pairs capable of forming overlaps. To determine whether our conclusions also hold at the genome-wide level, we extended the analysis to the complete dataset of closely spaced neighboring genes, without restricting it to the curated orthologous clusters. Specifically, we examined (i) whether overlap is associated with protein-complex membership according to STRING and (ii) whether overlap is associated with neighboring enzymatic reactions according to KEGG. Although this analysis is exploratory, because the annotation-cleaning procedure used for the orthologous clusters cannot be applied genome-wide, it provides an independent assessment of whether the observed patterns extend beyond the curated dataset.

The results for subunits were somewhat paradoxical. If we consider only pairs with an intergenic distance of at most 20 bp, overlapping genes form subunits even less frequently than non-overlapping ones, according to the Fisher exact test (odds ratio = 0.833). The same holds true, albeit to a lesser extent, for genes with an intergenic distance limit of 50 bp (odds ratio = 0.903). As the threshold is further increased, overlapping genes do indeed form subunits more often than non-overlapping ones. For metabolically related genes, overlap showed a positive association with metabolic linkage at both low intergenic-distance thresholds. At 20 bp, the effect was small (odds ratio = 1.114). At 50 bp, the association was stronger (odds ratio = 1.303), suggesting that metabolically linked neighboring genes are increasingly enriched for overlaps relative to non-linked pairs as the intergenic-distance threshold is relaxed.

## Discussion

In our study, the proportion of overlapping genes in unidirectional configuration turned out to be slightly higher (92.9%) than in previous studies (84%) [4]. This may be due to the fact that we collected statistics on the share of overlaps specifically for short overlaps. Similarly, the fact that most overlaps are either 1 or 4 bp in length has been observed previously [21]. We were able to confirm this with a larger sample than in previous studies.

Previously, analysis of genes orthologous to overlapping genes in *E. coli*, demonstrated that overlaps between two genes may be either lost or gained even among closely related species [9]. In our study, we confirmed this observation quantitatively. We therefore conclude that the evolution of overlapping genes is highly dynamic. Although these data should be interpreted with caution, as transition rate estimates may be less accurate for small clusters, the obtained values strongly support this conclusion.

No overall bias toward either overlap gain or loss was detected across the analyzed orthologous clusters. Instead, the estimates of transition rates exhibited substantial variation, with many clusters showing comparable gain and loss rates, while others were characterized by a pronounced excess of either gains or losses. Moreover, the estimated transition rates were generally high, indicating that overlap gain and loss can occur repeatedly over evolutionary time. The considerable heterogeneity observed among orthologous clusters suggests that the evolutionary dynamics of short gene overlaps are shaped by factors that vary between gene families, although the nature of these factors remains unclear. We also found no evidence that subunit-associated gene pairs exhibit systematically different overlap gain–loss dynamics. Thus, protein-complex association appears to affect neither the prevalence of overlap within orthologous clusters nor the relative balance between overlap gain and loss. We found no evidence that overlapping gene pairs are more likely to encode protein-complex subunits than gene pairs separated by short intergenic distances. In contrast, exploratory analyses even suggested a weak negative association between overlap and protein-complex membership at very short intergenic distances. These observations should nevertheless be interpreted with caution, as reliable overlap annotation is challenging on a genome-wide scale. In contrast, analyses based on orthologous clusters are likely to be more robust because overlap annotations within these clusters were manually curated and filtered to remove likely annotation artifacts. Taken together, our results indicate that gene overlap itself does not confer an increased tendency for genes to encode interacting protein-complex subunits beyond the effect of close genomic proximity.

This conclusion is consistent with previous studies. Earlier work showed that gene pairs that either overlap or are separated by short intergenic distances are more likely to encode protein-complex subunits than more distant gene pairs [11]. However, to the best of our knowledge, overlapping gene pairs have not previously been evaluated separately from other closely spaced gene pairs.

Interestingly, the genome-wide analysis revealed a modest positive association between overlap and metabolic linkage. The magnitude of this association increased as the distance threshold used to define non-overlapping neighbors was relaxed. However, this threshold dependence complicates interpretation, because larger thresholds introduce progressively more distant gene pairs into the comparison group. One possible explanation is that very closely spaced non-overlapping genes, like overlapping genes, are frequently located within the same operon and therefore already enriched for functional relationships. As the distance threshold increases, the comparison group is expected to contain a larger proportion of unrelated gene pairs not belonging to the same operon, potentially increasing the apparent enrichment associated with overlap. This interpretation is consistent with the known relationship between genomic proximity and metabolic organization [22]

This observation contrasts with the absence of a comparable enrichment for protein-complex subunits and may indicate that overlap is more strongly associated with the coordinated organization of metabolic pathways than with physical interactions between protein products. This interpretation should, however, be treated with caution, as our functional assignments provide only an approximate representation of the underlying biological relationships. In particular, the association between genes encoding interacting protein subunits may depend primarily on close genomic proximity, leading to spatial proximity of synthesized protein molecules, rather than on overlap per se. Moreover, given the large-scale nature of our analysis, we cannot exclude the possibility that overlap is important for the coordinated expression of protein-complex subunits in particular gene pairs or under specific biological conditions, even if such effects are not detectable as a general genome-wide trend.

## Declarations

### Ethics approval and consent to participate

Not applicable

### Consent for publication

Not applicable

### Availability of data and materials

The processed data supporting the findings of this study are openly available in Zenodo at https://zenodo.org/records/22093024 [23]. The repository includes genome-wide overlap data, orthologous cluster data, BayesTraits rate estimates, functional association datasets, and source data for representative ancestral-state reconstruction figures.

### Author Contributions

Conceptualization, A.S. and M.G.; methodology, A.S, A.K., and M.G.; software, A.S.; validation, A.S. and M.G; formal analysis, A.S.; investigation, A.S, A.K., and M.G.; resources, A.S and M.G..; data curation, A.S.; writing—original draft preparation, A.S.; writing—review and editing, A.K. and M.G.; visualization, A.S.; supervision, M.G.; project administration, M.G.; funding acquisition, M.G. All authors have read and agreed to the published version of the manuscript.

## Acknowledgements

Not applicable

